# Intra-peritoneal transplantation for efficient and easy generation of experimental acute myeloid leukemia in mice

**DOI:** 10.1101/2022.08.03.502696

**Authors:** Fenghua Qian, Brooke E. Arner, Shaneice K. Nettleford, Robert F. Paulson, K. Sandeep Prabhu

**Author notes:** **Corresponding author** (KSP). **Contributions:** K.S.P., R.F.P., and F.Q. contributed to study design, data interpretation, and wrote the manuscript; F.Q., B.E.A., and S.K.N. conducted experiments and data analyses.

## Abstract

**Objective:** There is an unmet need for novel therapies to treat acute myeloid leukemia (AML) and the associated relapse that involves leukemia stem cells (LSCs). An experimental AML rodent model to test therapies based on successfully transplanting these cells via retro-orbital injections in recipient mice is fraught with challenges. The aim of this study was to develop an easy, reliable, and consistent method to generate a robust murine model of AML using an intra-peritoneal route.

**Methods:** Bone marrow cells were transduced with a retrovirus expressing human MLL-AF9 fusion oncoprotein. Efficiency of lineage negative (Lin^−^) and Lin^−^Sca-1^+^c-Kit^+^ (LSK) populations as donor LSCs in the development of primary AML was tested. We developed a new method to generate AML by intra-peritoneal injection and discuss the pros and cons of intra-peritoneal and retro-orbital injections in serial transplantations.

**Results:** Both Lin^−^ and LSK cells transduced with human MLL-AF9 virus engrafted well in the bone marrow and spleen of recipients, leading to a full-blown AML. Intra-peritoneal injection of donor cells established AML in recipients upon serial transplantation.Infiltration of AML cells was detected in the blood, bone marrow, spleen, and liver of recipients by flow cytometry, qPCR, and histological analyses.

**Conclusions:**

1. Intra-peritoneal injection is an efficient method of AML induction using serial transplantation of donor cells.
2. Delayed homing to bone marrow, seen in intra-peritoneal injection that impedes the progression of AML, can be easily overcome by increasing the number of transplanted donor cells.
3. Increasing the number of transplanted donor cells can also help shorten the duration of onset of AML.
4. Sorting of LSK population within donor cells is not necessary in the generation of MLL-AF9-induced primary AML given that Lin^−^ population are equally efficient as LSK donor cells in primary transplantation of AML in mice.

## Introduction

Acute myeloid leukemia (AML) is a type of hematologic malignancy of diverse etiology with poor prognosis [1]. The generation of AML animal models lays the foundation for the understanding of its complex variations and pathobiology in an effort to discover novel therapies [2]. Leukemogenesis in mice involves transplantation of donor cells expressing fusion oncoproteins, including fusions involving the mixed lineage leukemia (MLL) gene to potently induce AML, to mimic the disease in humans [3]. Various cellular origins of donor cells have been reported in the transplantation of MLL gene-associated AML [4], with very little known about the cells responsible for the disease origin.

Multiple routes have been developed for transplantation in mice. Rather than intro-femoral injection, which directly introduces mutant donor cells into bone marrow [5], intravenous injection has been widely used to generate murine AML models that utilizes venous sinus plexus, tail vein, and jugular vein [6-9]. In the case of retro-orbital (r.o.) injection, various inherent disadvantages such as volume limitation, high technical demand, few chances for repeated attempt or error, and potential ocular injuries have been major stumbling blocks with limited or no viable alternatives. Tail vein injection can have similar problems besides local injuries. To facilitate the procedure, mice often need to be warmed up to dilate their tail veins [10]. Challenges include, hard to locate the tail vein without an additional light source, particularly in C57BL/6 strain of mice. For jugular vein injection, sufficient training of research personnel is necessary to locate the vein and limit possible complications. Besides, both venous sinus and jugular vein injections need to be performed under anesthesia, which adds another level of complexity. Thus, it is tempting to explore new routes for transplantation to facilitate the establishment of AML murine models.

Intra-peritoneal (i.p.) injection is commonly used to administer drugs, dyes, and anesthetics [11-15]. I.p. injection has been used to introduce hematopoietic cells for ectopic hematopoiesis [16] and transplant bone marrow-derived mesenchymal stem cells in various mouse models [17-21]. However, it has been infrequently used to establish hematopoietic malignancies in mice, particularly to study AML disease progression.

In the current study, we also describe the feasibility of i.p. injection in the generation of AML mouse models in addition to comparing the efficiency of donor cells from two common origins about AML transplantation. Our studies provide a simple and efficient way to generate experimental models of AML and related myeloid leukemias. Such a method has the potential to further our understanding of the disease mechanisms as well as provide us a relatively easy model to test experimental therapies.

## Materials and methods

### Plasmid transformation and transfection

α-Select competent cells (BIO-85027, Bioline) were transformed with MSCV-MLL-AF9-EF1α-luc2-P2A-EGFP-LC3 plasmids. Antibiotic (ampicillin) selection was applied to isolate transformed cells that were used to isolate plasmids in large scale. Plasmids were extracted using QIAGEN Plasmid Maxi Kit according to manufacturer’s instructions. To generate retrovirus, Phoenix Ecotropic (pECO) cells (ATCC CRL-3214) were transfected with plasmids using *Trans*IT-293 transfection reagent (MIR2705, Mirus). Briefly, pECO cells were seeded in 6 cm^2^ dishes and transfection was initiated when cells were at 50∼60% confluence with 5.5 μg MSCV-MLL-AF9 plasmids and 16.4 μL *Trans*IT-293 transfection reagent according to the manufacturer’s instructions. Supernatants were collected and passed through a 0.45 μm filter after 48 h and stored in -80 °C.

### Transduction and transplantation

Eight-to ten-week-old male C57BL6/J mice (CD45.2) were purchased from Taconic Biosciences. Eight-to twelve-week-old CD45.1 donor mice were euthanized, and bone marrow cells were flushed from femurs and tibias followed by lysis of red blood cell (RBC) with ACK lysis buffer (155 mM NH_4_Cl, 12 mM KHCO_3_, 0.1mM EDTA-2Na).Lineage negative (Lin^−^) cells were enriched using EasySep™ Mouse Hematopoietic Progenitor Cell Isolation Kit (StemCell technologies) and stained with fluorescent antibodies: APC-conjugated anti-mouse CD117 (c-Kit) (Biolegend) and PE-Cy7-conjugated anti-mouse Ly-6A/E (Sca-1) (BD Biosciences). HSCs were further gated as Lin^−^Sca-1^+^c-Kit^+^ (LSK) population in a Beckman Coulter MoFlo Astrios EQ Cell Sorter [22, 23]. Unsorted Lin^−^ cells or sorted HSCs were cultured in a retronectin-coated 6 cm^2^ dish with 3 mL viral supernatants for 6 h in 3 mL IMDM media supplemented with 15% heat inactivated fetal bovine serum (Atlanta Biologicals), 1% bovine serum albumin (Gemini bio-products), 10 mg/mL insulin (Sigma), 20 mg/mL Holo transferrin (Sigma Aldrich), 1% L-glutamine (HyClone), 10 mg/mL ciprofloxacin (GoldBio), 0.007‰ (v/v) β-mercaptoethanol (Sigma Aldrich), 25 ng/mL mouse recombinant (mr) stem cell factor (Gemini bio-products), 25 ng/mL mrFlt3L (Pepro Tech), 10 ng/mL mrIL-3 (Gemini bio-products), and 10 ng/mL mrIL-6 (Gemini bio-products). Cells were harvested, resuspended in PBS, and injected retro-orbitally (0.1 mL PBS/mouse, Lin^−^ cells or LSK cells) or intra-peritoneally (0.5 mL PBS/mouse, Lin^−^ cells) into CD45.2 male recipient mice (Fig 1).

**Fig 1.**
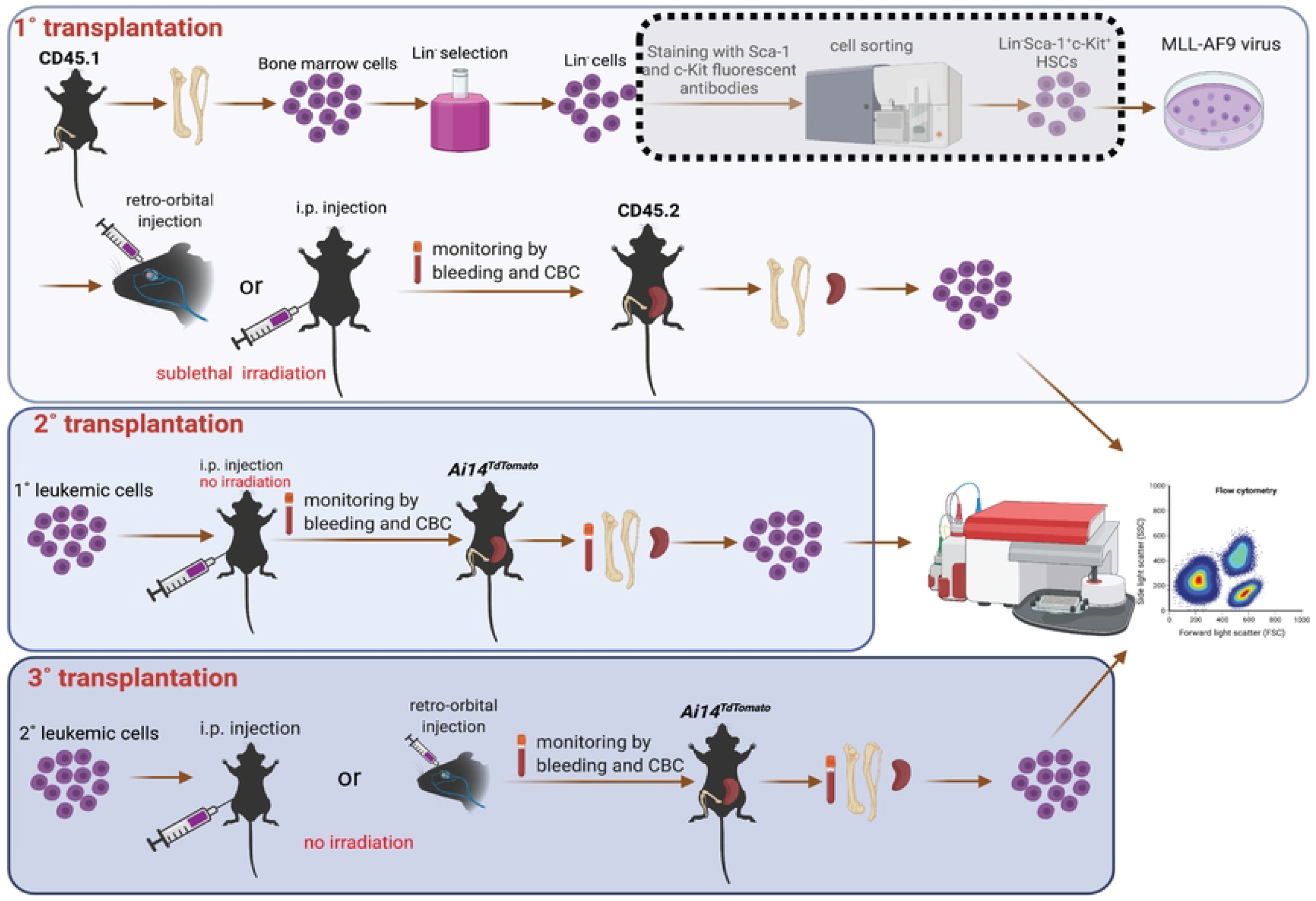
Schematic of MLL-AF9 viral transduction in bone marrow HSCs and serial transplantation (1°, 2°, and 3°). Sorting of Sca-1 and c-Kit double positive population by Beckman Coulter MoFlo Astrios EQ Cell Sorter shown in the dotted shade box is considered optional, should resources allow. The figure was created using BioRender (https://biorender.com/).

For primary (1°) transplantation, recipient mice were sublethally irradiated (4.75 Gy) to facilitate engraftment. Mice were provided water *ad libitum* containing antibiotics to prevent opportunistic digestive infections from three days prior to transplantation to seven days post transplantation. AML was initially diagnosed by leukocytosis and further confirmed by hepatosplenomegaly and infiltration of AML cells in the bone marrow and spleen. Furthermore, mice were monitored for leukocytosis by evaluating complete blood count (CBC) in retro-orbitally collected peripheral blood on a Hemavet 950FS (Drew Scientific). At the endpoint [white blood cell (WBC) > 40 K/μL], splenocytes were harvested by mashing spleens through a 70 μm sterile strainer, and bone marrow cells were isolated by flushing femurs and tibias followed by RBC lysis. AML cells were identified by staining total splenocytes and bone marrow cells with FITC-conjugated anti-mouse CD45.1 antibody on a BD Accuri C6 flow cytometer (Fig 1).

For secondary (2°) transplantation, AML cells from 1° i.p. recipients were resuspended in PBS (0.5 mL/mouse) and injected intra-peritoneally into non-irradiated red fluorescence protein (RFP)-expressing *Ai14*^*TdTomato*^ male mice without antibiotics-containing water. In a parallel experiment, CD45.1 AML cells in PBS (0.1 mL/mouse) from 1° r.o. recipients were injected retro-orbitally into CD45.2 male mice without irradiation or antibiotic water. By monitoring the CBC in peripheral blood, we identified the disease progression in 2° recipients. To further confirm the establishment of AML, we collected whole peripheral blood by heart puncture as well as bone marrow, spleen, and liver. Furthermore, we performed i.p. lavage to collect i.p. cells. Single cell suspensions were acquired from bone marrow, spleen, and i.p. lavage as described above. Cells from blood, bone marrow, spleen, and peritoneal cavity were analyzed on a BD Accuri C6 flow cytometer following RBC lysis. AML cells were recognized as RFP negative (RFP^−^) cells.

For tertiary (3°) transplantation, *Ai14*^*TdTomato*^ (RFP^+^) or CD45.2 mice were intra-peritoneally injected with AML cells from bone marrow or peritoneal cavity of 2° i.p. recipients, respectively. R.o. injection was done on *Ai14*^*TdTomato*^ (RFP^+^) mice with AML cells from peritoneal cavity of 2° r.o. recipients. At the endpoint, we sampled blood, bone marrow, spleen, liver, and i.p. cells; RFP^−^ or CD45.1^+^ cells were also examined by flow cytometry.

For i.p. lavage, 5 mL IMDM media/time was injected into the peritoneal cavity twice to collect the cells. Cell numbers for transplantations are described in detail in figure legends. All experiments described above were preapproved by the Institutional Animal Care and Use Committee (IACUC) at The Pennsylvania State University.

### Histological analysis

Isolated spleen, liver, and femur of mice were isolated upon euthanasia and fixed in 10% (v/v) buffered formalin. Spleen and liver from a healthy counterpart were also sampled for comparisons. The fixed tissues were embedded by paraffin and cut into sections. Subsequently, sections were stained with hematoxylin and eosin (H&E) dyes. Images were obtained under a microscope installed with AxioVision software for histological analysis.

### Real-time PCR

RNAs were prepared in TRI reagent® solution (Sigma) according to manufacturer’s instructions. RNAs were used to synthesize cDNA by a High-Capacity cDNA Reverse Transcription Kit (Applied biosystems). cDNA was applied to perform qPCR using PerfeCTa qPCR SuperMix (Quanta Biosciences) in a 7300 Real time PCR system (Applied Biosystems). The following pre-validated TaqMan probes were used: *KMT2A* (*MLL*) (Ref Seq: NM_001197104(2), IDT) [24] and *18S* ribosomal RNA (Hs99999901_s1). *KMT2A* and *18S* amplicons were loaded onto an agarose gel to visualize the expression. Images were acquired in an imager (G:Box Chemi) installed with GeneSys software program (Version 1.5.7.0).

### Data processing

Results were analyzed using GraphPad Prism version 6 (GraphPad) and presented as the mean ± SEM. Figures were generated using Adobe illustrator (Version 25.2.3).

## Results

### Comparison of the transplantation efficiency of murine AML cells using r.o. and i.p. routes of transplantation

We previously reported the establishment of 1° AML in recipient mice retro-orbitally transplanted with MLL-AF9-transduced LSK cells and demonstrated the transplantable of 1° AML cells by serial transplantation [25]. Here, we first evaluated the possibility of using bone marrow Lin^−^ cells to perform transplantation. The presence of aberrant leukocytosis (Fig 2A) and increased infiltration of leukemic cells (CD45.1^+^) in the bone marrow and spleen (Fig 2B) supported the feasibility of using bone marrow Lin^−^ cells to generate 1° AML. To compare the disease induction period, two donor mice per recipient were euthanized to isolate either LSK or Lin^−^ cells, no significant differences were observed in 1° recipients transplanted with LSK or Lin^−^ cells (Fig 2C).

**Fig 2.**
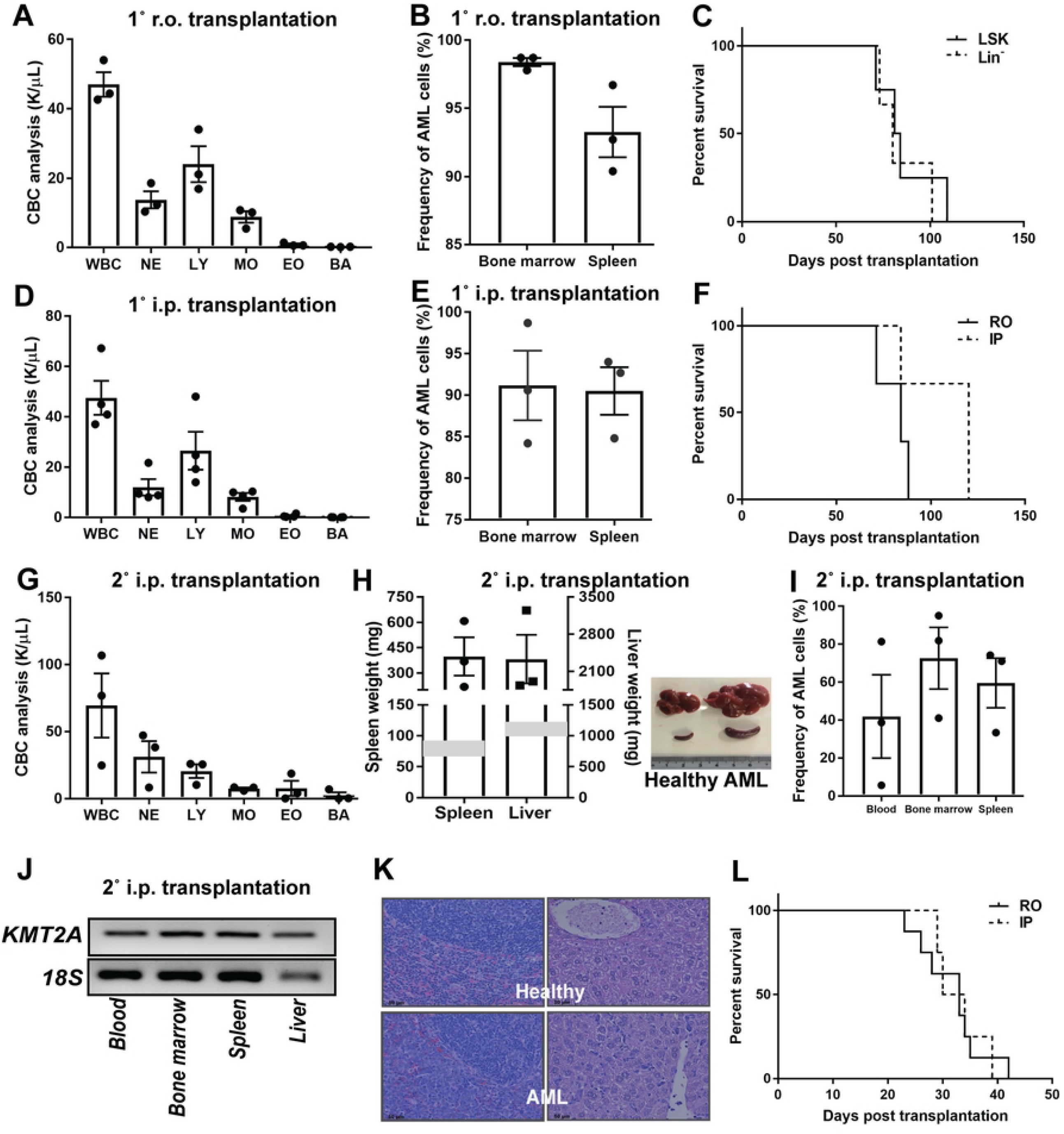
Generation of murine AML model in 1° and 2° transplantation. A. CBC (K/μL blood), WBC, neutrophil (NE), lymphocyte (LY), monocyte (MO), eosinophil (EO), and basophil (BA) profile of 1° recipient mice transplanted retro-orbitally with MLL-AF9-transduced bone marrow Lin^−^ cells (5.125 × 10^6^ cells/mouse) by hemavet (n=3). B. Flow cytometric analysis of the frequency of AML cells (CD45.1^+^) in the bone marrow and spleen of 1° CD45.2 recipient mice transplanted retro-orbitally with MLL-AF9-transduced bone marrow Lin^−^ (5.125 × 10^6^ cells/mouse) cells (n=3). C. Incubation duration of 1° AML in recipients transplanted retro-orbitally with MLL-AF9-transduced bone marrow LSK cells (3.16 × 10^5^/mouse isolated from two donors, n=4) or Lin^−^ cells (3.69 × 10^6^/mouse isolated from two donors, n=3). Each recipient received cells isolated from two donors. D. CBC analysis of 1° recipient mice transplanted intra-peritoneally with MLL-AF9-transduced bone marrow Lin^−^ cells (5.125 × 10^6^ cells/mouse) by hemavet (n=4). E. Flow cytometric analysis of the frequency of AML cells (CD45.1^+^) in the bone marrow and spleen of 1° CD45.2 recipient mice transplanted intra-peritoneally with MLL-AF9-transduced bone marrow Lin^−^ cells (5.125 × 10^6^ cells/mouse, n=3). F. Incubation duration of 1° AML in recipients transplanted retro-orbitally (n=3) or intra-peritoneally (n=3) with MLL-AF9-transduced bone marrow Lin^−^ cells. Each recipient received same amount of donor cells (5.125 × 10^6^/mouse). G. CBC analysis of 2° recipients transplanted intra-peritoneally with AML cells (8 × 10^5^ cells/mouse) isolated from the spleen of 1° i.p. transplanted recipients by hemavet (n=3). H. Weights (mg) of spleen and liver of 2° recipient mice transplanted intra-peritoneally with AML cells (8 × 10^5^ cells/mouse) isolated from the spleen of 1° i.p. transplanted recipients. The shade areas represent the normal weight ranges of spleen and liver. Representative image of spleen and liver from 2° recipient mice and healthy counterparts. I. Flow cytometric analysis of the frequency of AML cells (RFP^−^) in the blood, bone marrow, and spleen of 2° RFP^+^ recipient mice transplanted intra-peritoneally with AML cells (8 × 10^5^ cells/mouse) isolated from the spleen of 1° i.p. transplanted recipients (n=3). J. Gene expression of *KMT2A* and *18S* presented as DNA band. RNAs were isolated from the blood, bone marrow, spleen, and liver of 2° recipient mice transplanted intra-peritoneally with AML cells (8 × 10^5^ cells/mouse) isolated from the spleen of 1° i.p. transplanted recipients. K. Representative images of histological changes in the spleen (left panel) and liver (right panel) of 2° recipients transplanted intra-peritoneally with AML cells (8 × 10^5^ cells/mouse) isolated from the spleen of 1° i.p. transplanted recipients and healthy counterparts. L. Incubation duration of 2° AML in recipients transplanted retro-orbitally (4 × 10^5^/mouse, n=9) or intra-peritoneally (8 × 10^5^/mouse, n=4) with AML cells isolated from 1° r.o. and i.p. transplanted recipients, respectively.

During the examination of metastasis of 1° AML, we surprisingly discovered the spread of AML cells in abdominal cavity of 1° recipients with MLL-AF9-transduced Lin^−^ cells (Fig S1). This led to us to explore if i.p. injection could be adopted in the generation of murine AML, which might serve as a new and easier route of transplantation. Due to the success of using bone marrow Lin^−^ population as AML donor cells, we confirmed establishment of 1° AML via i.p. injection of MLL-AF9-transduced bone marrow Lin^−^ cells, as seen in the form of leukocytosis (Fig 2D) and presence of AML cells (CD45.1^+^) in the bone marrow and spleen of recipient mice (Fig 2E).However, transplantation via i.p. injection took longer time to develop AML than via r.o. injection despite the recipients receiving the equal number of donor cells (Fig 2F).Besides, mice transplanted retro-orbitally with more Lin^−^ cells in Fig 2F seemed to develop AML faster than those transplanted with fewer Lin^−^ cells in Fig 2C.

Owing to the inherent inconsistency of incubation time in 1° AML recipients (Fig 2F), we sought to test the i.p. transplantation in 2° recipients. For 2° transplantation, i.p. injection was performed with 8 × 10^5^ AML cells isolated from intra-peritoneally transplanted 1° recipients. 2° recipients showed leukocytosis (Fig 2G) and significant hepatosplenomegaly (Fig 2H) in less than one month post transplantation, a clear advancement from 1° transplantation. Consistent with this observation, we also detected AML cells (RFP^−^) in the peripheral blood, bone marrow, and spleen (Fig 2I) as well as in the peritoneal cavity (Fig S1). Besides, the expression of oncogene *KMT2A* in the blood, spleen, and liver of 2° recipients also confirmed the successful generation of AML (Fig 2J). Moreover, histological analysis further demonstrated the infiltration of leukemic cells in the spleen and liver of intra-peritoneally transplanted mice (Fig 2K). These data confirm that i.p. injection-generated 1° AML cells are transplantable in 2° recipients.Furthermore, i.p. injection of 8 × 10^5^ AML cells per mouse into recipients achieved comparable engraftment to r.o. injection of 8 × 10^5^ AML cells per mouse into recipients (Fig 2L).

One of the key features of LSCs is serial transplantability. Thus, to further ascertain the establishment of AML in 2° recipients, we attempted to see if AML cells from 2° i.p. transplantation remained serially transplantable. Meanwhile, we also attempted to verify the feasibility of stem cell transplantation by i.p. injection in 3° transplantation. AML cells isolated from either bone marrow (8 × 10^5^ leukemic cells/mouse in 0.5 mL PBS) or i.p. lavage (4 × 10^5^ leukemic cells/mouse in 0.5 mL PBS) of 2° recipients were injected into 3° recipients. Although being transplanted with donor cells generated from different sources, 3° recipient mice acutely exhibited AML signs (three and four weeks for bone marrow cells and i.p. cells, respectively), including leukocytosis (Fig 3A) and hepatosplenomegaly (Fig 3B), presence of leukemic cells in the blood, bone marrow, spleen (Fig 3C), and expression of *KMT2A* (Fig 3D). Histological observation of femur and spleen further demonstrated the infiltration of leukemic cells (Fig 3E). As a comparison, r.o. injection of leukemic splenocytes (4 × 10^5^ cells) from 2° recipients was also equally capable of engrafting and developing AML in 3° recipients, characterized by leukocytosis (Fig S2A), hepatosplenomegaly (Fig S2B), and infiltration of AML cells in the blood, bone marrow, and spleen (Fig S2C).

**Fig 3.**
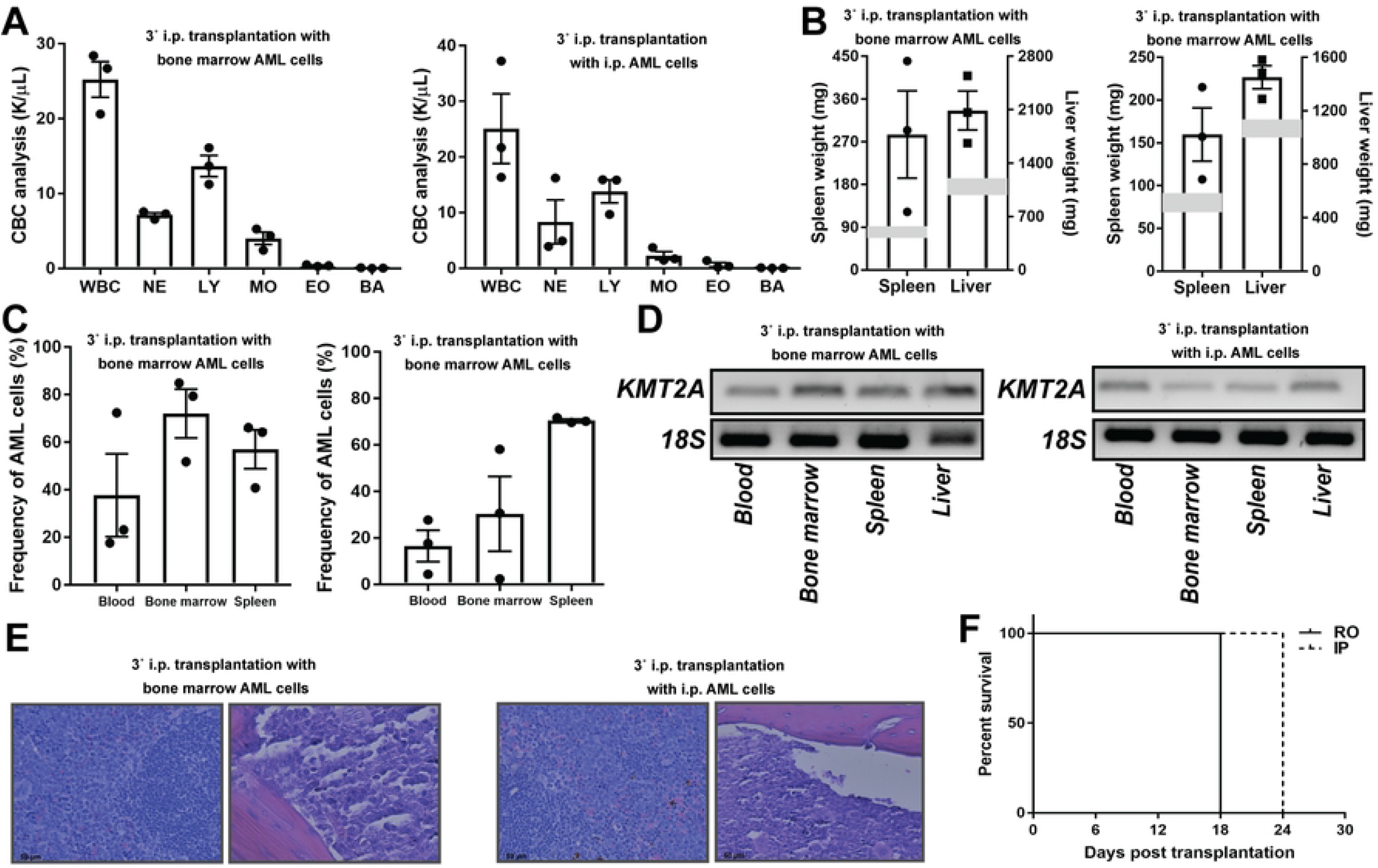
3° transplantation of AML cells via i.p. transplantation. (A-E) 3° RFP^+^ or CD45.2 recipient mice. Left: i.p. injection of AML cells (8 × 10^5^/mouse) from bone marrow of 2° recipients; Right: i.p. injection of AML cells (4 × 10^5^/mouse) from peritoneal cavity (i.p.) of 2° recipients. n=3. A. CBC analysis of 3° recipient mice by hemavet. B. Weights (mg) of spleen and liver of 3° recipient mice. The shade areas represent the normal weight ranges of spleen and liver. C. Flow cytometric analysis of the frequency of AML cells (left: RFP^−^; right: CD45.1^+^) in the blood, bone marrow, and spleen of 3° recipient mice. D. Gene expression of *KMT2A* and *18S* presented as DNA band. RNAs were isolated from the blood, bone marrow, spleen, and liver of 3° recipient mice. E. Representative images of histological changes in the spleen (left panel) and femur (right panel) of 3° recipient mice. F. Incubation duration of 3° AML in recipients transplanted retro-orbitally (4 × 10^5^/mouse, n=3) or intra-peritoneally (4 × 10^5^/mouse, n=3) with AML cells isolated from 2° r.o. and i.p. transplanted recipients, respectively.

### Intra-peritoneal transplantation is effective even for 3° transplantation of AML cells

Compared to the 2° and 3° transplantations, we could easily conclude that 3° transplantation (Fig 3F) progressed much faster than 2° transplantation (Fig 2L) for both i.p. and r.o. injections. Notably, i.p. injection of 4 × 10^5^ AML cells/mouse developed AML later than r.o. injection of same number of AML cells for 6 days (Fig 3F), possibly indicating the time spent on the migration from peritoneal cavity to bone marrow niche. Interestingly, the high frequency of leukemic cells in the peritoneal cavity (Fig S1) in the 3° transplanted mice suggested that peritoneal cavity may also provide the niche for the fast expansion of leukemic cells. Besides, the presence of leukemic cells in the peritoneal cavity of 1° and 3° recipients transplanted retro-orbitally indicated the mutual circulation of leukemic cells between peritoneal cavity and blood system, justifying the i.p. transplantation. Collectively, these findings validate the use of i.p. injection in serial transplantation of AML.

## Discussion

These preliminary findings provide supportive evidence that transplantation of Lin^−^ cells is comparable to LSK cells in the generation of 1° murine AML. In addition, our data also showed that i.p. injection is an efficient and convenient method to establish murine AML compared to intravenous (or r.o.) injection.

In addition to LSK cells, other populations such as granulocyte-monocyte progenitors, common lymphoid progenitors, and common myeloid progenitors have been reported to be substituted as donor cells in the generation of 1° MLL-AF9-induced AML with various incubation durations [26, 27]. Quantitative comparison is lacking to suggest the optimal choice. In the current study, we showed that Lin^−^ and LSK isolated from same donor mice had no apparent difference in the generation of 1° MLL-AF9-induced AML, given that transduced myeloid progenitor cells (Sca-1^−^) were able to initiate AML [28].Considering the expense and time required for flow cytometry-based sorting of LSK population, these studies support the use of Lin^−^ population as the donor cells for 1° transplantation.

In terms of the amount of donor mice in 1° transplantation, one donor was able to provide ∼1.0 to 1.5 × 10^5^ LSK cells, and one recipient transplanted with this amount of LSK cells developed 1° AML in about four months. However, increasing the number of donors to build up LSK cells up to ∼3 × 10^5^ per recipient can shorten the incubation duration to ∼70 days. Thus, two donors should be enough to rapidly generate 1° AML. This principle also applies to Lin^−^ cells wherein ∼1.0 to 2.5 × 10^6^ Lin^−^ cells can be isolated from each donor mouse. However, when donor cell density was increased, the turnaround time of leukemogenesis was considerably shorter.

Compared to r.o. injection, i.p. injection took about ∼20 more days to develop the full-blown 1° AML, whereas this limitation was overcome by increasing the donor cells for i.p. injection. Despite such an apparent difference, both methods showed similar engraftment rate (>80%) in the bone marrow and spleen at the endpoint, indicating the general applicability of i.p. injection in 1° transplantation in AML.

The relatively long incubation period for 1° transplantation as well as its unpredictable disease progression in terms of inconsistency of latency in recipient mice is a major limitation for controlled experiments, such as pharmacological interventions and mechanistic studies. Thus, we recommend using 1° leukemic cells for transplantation in subsequent serial transplantations, typically 2° transplantation (Fig. 1). Given that up to 3 × 10^8^ leukemic cells in spleen (and 6 × 10^7^ in bone marrow) can be generated from every single 1° recipient at the end point, which would be sufficient to conduct 2° transplantation on more than three hundred mice, we suggest transplanting as many cells as possible to fewer recipients to shorten the latency and increase engraftment rate of 1° transplantation. Given its consistent and short latency, 2° transplantation can be utilized in multiple applications, including *in vivo* experiments mentioned above.Furthermore, i.p. injection also facilitates the procedure for abundant transplantation for inexperienced technicians. Compared to r.o. injection route, which generally generates the AML in three weeks, i.p. injection method took relatively longer (∼ four weeks) with the same amount of donor cells to establish a full-blown AML. Even though it is relatively easier and more convenient, increasing the transplanted cell number or performing tertiary transplantation can also accelerate the progression of AML by i.p. injection.

## Acknowledgments

The authors thank Huck Institute’s Flow Cytometry Core Facility, and Histopathology Core Facility of the Animal Diagnostic Laboratory, Department of Veterinary and Biomedical Sciences, The Pennsylvania State University, for providing timely technical support. This work was supported by grants from American Institute for Cancer Research (KSP), Penn State College of Agricultural Sciences, Penn State Cancer Institute, USDA-NIFA project 4771, Accession number 00000005 to KSP and RFP.

## Supporting information

**S1 Fig. Flow cytometric analysis of AML cell infiltration in the peritoneal cavity**. Frequency of AML cells in the peritoneal lavage of 1° recipients transplanted retro-orbitally (1° RO, n=2), 2° recipients transplanted intra-peritoneally (2° IP, n=2), and 3° recipients transplanted via i.p. injection with bone marrow AML cells (3° BM-IP, n=3) and i.p. AML cells (3° IP-IP, n=3), as well as 3° recipients transplanted via r.o. injection of AML splenocytes (3° SP-RO, n=3).

**S2 Fig. 3° transplantation of AML splenocytes via r.o. injection**.A. CBC analysis of 3° recipient mice retro-orbitally transplanted with AML cells (4 × 10^5^/mouse) from spleen of 2° recipient mice. B. Weights (mg) of spleen and liver of 3° retro-orbitally transplanted recipient mice. The shade areas represent the normal weight ranges of spleen and liver. C. Frequency of AML cells (RFP^−^) in the blood, bone marrow, and spleen of 3° retro-orbitally transplanted recipient mice (n=3).

## Notes

### Competing Interest Statement

The authors have declared no competing interest.

## References

1. Dohner K, Paschka P, Dohner H. Acute myeloid leukemia. Internist. 2015;56(4):354–63. doi: 10.1007/s00108-014-3596-5. PubMed PMID: WOS:000352643200004.

2. Fortier JM, Graubert TA. Murine models of human acute myeloid leukemia. Cancer Treat Res. 2010;145:183-96. Epub 2009/01/01. doi: 10.1007/978-0-387-69259-3_11. PubMed PMID: 20306252.

3. Ernst P, Fisher JK, Avery W, Wade S, Foy D, Korsmeyer SJ. Definitive hematopoiesis requires the mixed-lineage leukemia gene. Dev Cell. 2004;6(3):437–43. doi: Doi 10.1016/S1534-5807(04)00061-9. PubMed PMID: WOS:000222442900016.

4. Fisher JN, Kalleda N, Stavropoulou V, Schwaller J. The Impact of the Cellular Origin in Acute Myeloid Leukemia: Learning From Mouse Models. Hemasphere. 2019;3(1). PubMed PMID: WOS:000507915600001.

5. Zhan Y, Zhao Y. Hematopoietic stem cell transplant in mice by intra-femoral injection. Methods Mol Biol. 2008;430:161-9. Epub 2008/03/29. doi: 10.1007/978-1-59745-182-6_11. PubMed PMID: 18370298.

6. Price JE, Barth RF, Johnson CW, Staubus AE. Injection of cells and monoclonal antibodies into mice: comparison of tail vein and retroorbital routes. Proceedings of the Society for Experimental Biology and Medicine. 1984;177(2):347–53.

7. Yardeni T, Eckhaus M, Morris HD, Huizing M, Hoogstraten-Miller S. Retro-orbital injections in mice. Lab Anim (NY). 2011;40(5):155–60. Epub 2011/04/22. doi: 10.1038/laban0511-155. PubMed PMID: 21508954; PubMed Central PMCID: PMCPMC3158461.

8. Suckow MA, Danneman P, Brayton C. The laboratory mouse: CRC Press Inc.; 2001.

9. Barr JE, Holmes DB, Ryan LJ, Sharpless SK. Techniques for the chronic cannulation of the jugular vein in mice. Pharmacol Biochem Behav. 1979;11(1):115–8. Epub 1979/07/01. doi: 10.1016/0091-3057(79)90307-1. PubMed PMID: 493295.

10. Kang Y. Analysis of cancer stem cell metastasis in xenograft animal models. Cancer stem cells: Springer; 2009. p. 7–19.

11. Nungestee W, Wolf A, Jourdonais L. Effect of gastric mucin on virulence of bacteria in intraperitoneal injections in the mouse. Proceedings of the Society for Experimental Biology and Medicine. 1932;30(2):120–1.

12. Gargiulo S, Greco A, Gramanzini M, Esposito S, Affuso A, Brunetti A, et al. Mice anesthesia, analgesia, and care, Part I: anesthetic considerations in preclinical research. ILAR journal. 2012;53(1):E55–E69.

13. Leong S-K, Ling E-A. Labelling neurons with fluorescent dyes administered via intravenous, subcutaneous or intraperitoneal route. Journal of neuroscience methods. 1990;32(1):15–23.

14. Ma P, Luo Q, Chen J, Gan Y, Du J, Ding S, et al. Intraperitoneal injection of magnetic Fe3O4-nanoparticle induces hepatic and renal tissue injury via oxidative stress in mice. International journal of nanomedicine. 2012;7:4809.

15. Schwarze SR, Ho A, Vocero-Akbani A, Dowdy SF. In vivo protein transduction: delivery of a biologically active protein into the mouse. Science. 1999;285(5433):1569–72.

16. Muench MO, Chen JC, Beyer AI, Fomin ME. Cellular therapies supplement: the peritoneum as an ectopic site of hematopoiesis following in utero transplantation. Transfusion. 2011;51:106s–17. PubMed PMID: WOS:000297290700006.

17. Zhao W, Li J-J, Cao D-Y, Li X, Zhang L-Y, He Y, et al. Intravenous injection of mesenchymal stem cells is effective in treating liver fibrosis. World journal of gastroenterology: WJG. 2012;18(10):1048.

18. Yousefi F, Ebtekar M, Soleimani M, Soudi S, Hashemi SM. Comparison of in vivo immunomodulatory effects of intravenous and intraperitoneal administration of adipose-tissue mesenchymal stem cells in experimental autoimmune encephalomyelitis (EAE). Int Immunopharmacol. 2013;17(3):608–16. Epub 2013/08/27. doi: 10.1016/j.intimp.2013.07.016. PubMed PMID: 23973288.

19. Cheng K, Rai P, Plagov A, Lan XQ, Kumar D, Salhan D, et al. Transplantation of bone marrow-derived MSCs improves cisplatinum-induced renal injury through paracrine mechanisms. Exp Mol Pathol. 2013;94(3):466–73. PubMed PMID: WOS:000319535900007.

20. Castelo-Branco MT, Soares ID, Lopes DV, Buongusto F, Martinusso CA, do Rosario A, Jr., et al. Intraperitoneal but not intravenous cryopreserved mesenchymal stromal cells home to the inflamed colon and ameliorate experimental colitis. PLoS One. 2012;7(3):e33360. Epub 2012/03/21. doi: 10.1371/journal.pone.0033360. PubMed PMID: 22432015; PubMed Central PMCID: PMCPMC3303821.

21. Bazhanov N, Ylostalo JH, Bartosh TJ, Tiblow A, Mohammadipoor A, Foskett A, et al. Intraperitoneally infused human mesenchymal stem cells form aggregates with mouse immune cells and attach to peritoneal organs. Stem Cell Res Ther. 2016;7:27. Epub 2016/02/13. doi: 10.1186/s13287-016-0284-5. PubMed PMID: 26864573; PubMed Central PMCID: PMCPMC4748482.

22. Wognum AW, Eaves AC, Thomas TE. Identification and isolation of hematopoietic stem cells. Arch Med Res. 2003;34(6):461–75. Epub 2004/01/22. doi: 10.1016/j.arcmed.2003.09.008. PubMed PMID: 14734086.

23. Randall TD, Weissman IL. Characterization of a population of cells in the bone marrow that phenotypically mimics hematopoietic stem cells: resting stem cells or mystery population? Stem Cells. 1998;16(1):38–48. Epub 1998/02/25. doi: 10.1002/stem.160038. PubMed PMID: 9474746.

24. Ronan JL, Wu W, Crabtree GR. From neural development to cognition: unexpected roles for chromatin. Nat Rev Genet. 2013;14(5):347–59. Epub 2013/04/10. doi: 10.1038/nrg3413. PubMed PMID: 23568486; PubMed Central PMCID: PMCPMC4010428.

25. Qian F, Arner BE, Kelly KM, Annageldiyev C, Sharma A, Claxton DF, et al. Interleukin-4 treatment reduces leukemia burden in acute myeloid leukemia. FASEB journal: official publication of the Federation of American Societies for Experimental Biology. 2022;36(5):e22328.

26. Krivtsov AV, Twomey D, Feng ZH, Stubbs MC, Wang YZ, Faber J, et al. Transformation from committed progenitor to leukaemia stem cell initiated by MLL-AF9. Nature. 2006;442(7104):818–22. PubMed PMID: WOS:000239792700047.

27. Chen W, Kumar AR, Hudson WA, Li Q, Wu B, Staggs RA, et al. Malignant transformation initiated by Mll-AF9: gene dosage and critical target cells. Cancer Cell. 2008;13(5):432–40. Epub 2008/05/06. doi: 10.1016/j.ccr.2008.03.005. PubMed PMID: 18455126; PubMed Central PMCID: PMCPMC2430522.

28. Somervaille TCP, Cleary ML. Identification and characterization of leukemia stem cells in murine MLL-AF9 acute myeloid leukemia. Cancer Cell. 2006;10(4):257–68. PubMed PMID: WOS:000241714400003.

